## Supplementary information for "Marcksl1 modulates endothelial cell mechanoresponse to haemodynamic forces to control blood vessel shape and size"

#### SUPPLEMENTARY FIGURE LEGENDS

**Supplementary Fig. 1. *marcksl1a* and *marcksl1b* mRNA expression**

**a**, *In situ* hybridization with *marcksl1a* and *marcksl1b* RNA probes at different developmental stages. The inset shows a magnified view of the caudal vein plexus (CVP). **b**, **c**, scRNAseq analysis showing *marcksl1a* and *marcksl1b* expression levels in ECs of 1 and 3 dpf embryos.

**Supplementary Fig. 2. Marcksl1 expression level regulates blood vessel diameter**

Plasmid constructs encoding *fli1ep*-driven expression of full-length (**a**) or mutated (**b**, without an Effector domain, ΔED) Marcksl1a or Marcksl1b proteins tagged with EGFP. Maximum intensity projection of a confocal z-stack of ISVs in zebrafish trunk of 2 dpf *Tg(fli1:myr-mCherry)*$^{ncv1}$ transgenic embryos expressing full length Marcksl1a (**c**), Marcksl1aΔED (**d**), full length Marcksl1b (**f**) or Marcksl1aΔED (**g**). Marcksl1 overexpressing cells are in green. **e,** Quantification of aISV, vISV and DLAV diameter in control and Marcksl1a-overexpressing blood vessels (control: $n$=27 aISVs/30 vISVs/24 DLAVs from 34 embryos; full length Marcksl1a: $n$=25 aISVs/18 vISVs/13 DLAVs from 20 embryos; Marcksl1aΔED: $n$=17 aISVs/13 vISVs/9 DLAVs from 14 embryos). **h**, Quantification of aISV, vISV and DLAV diameter in control and Marcksl1b-overexpressing vessels (control: $n$=17 aISVs/15 vISVs/21 DLAVs from 38 embryos; full length Marcksl1b: $n$=21 aISVs/18 vISVs/22 DLAVs from 23 embryos; Marcksl1aΔED: $n$=25 aISVs/10 vISVs/14 DLAVs from 15 embryos). Statistical significance was determined by ordinary one-way ANOVA with Tukey′s multiple comparisons test. $^{*}P < 0.05$, $^{**}P < 0.01$, $^{***}P < 0.001$, $^{****}P < 0.0001$. Scale bars, 20 μm.

**Supplementary Fig. 3. Mouse and zebrafish Marcksl1 proteins share sequence homology.**

The conserved positively charged Effector Domain and putative phosphorylation sites (in red), T124 in Marcksl1a and T162 in Marcksl1b, are shown. T124 and T162 sites correspond to experimentally validated[10] JNK kinase phosphorylation sites, S120 and T148, respectively, in mouse Marcksl1.

**Supplementary Fig. 4. Generation of *marcksl1a* and *marcksl1b* mutant zebrafish**

**a, b,** CRISPR/Cas9- and TALEN-mediated mutagenesis of *marcksl1a* and *marcksl1b* genes, respectively. **a**) Zebrafish *marcksl1a* gene structure, gRNA binding site (in red), *marcksl1a*$^{rk23}$ mutant allele, Marcksl1a wild type (213 aa) and Marcksl1a$^{rk23}$ (truncated at 56 aa) protein structure. The *rk23* mutation causes a 2-nt deletion which leads to a frameshift after 51 aa and premature stop codon (underlined), and as a result to the loss of the Effector Domain and JNK phosphorylation site, T124. **b,** Zebrafish *marcksl1b* gene structure, a pair of TALEN binding sites (in red), *marcksl1b*$^{rk24}$ mutant allele, Marcksl1b wild type (207 aa) and Marcksl1b$^{rk24}$ (truncated at 17 aa) protein structure. The *rk24* mutation causes a 5-nt deletion which leads to a frameshift after 13 aa and premature stop codon (underlined), and as a result to loss of MH2 and Effector domains and JNK phosphorylation site, T162. Non-coding and coding parts of exons are shown as white and dark blue rectangles, respectively. The partial nucleotide and amino acid sequences are shown. **c, d**, Sequence reads showing 2-nt and 5-nt deletions (shadowed area in wild type) in *marcksl1a*$^{rk23}$ (**c**) and *marcksl1b*$^{rk24}$ (**d**) alleles (red dotted line), respectively. Premature stop codons are underlined. Amino acid sequences are shown above the nucleotide sequences. **e**) Bright field images of wild type, *marcksl1a*$^{rk23}$, *marcksl1b*$^{rk24}$ and *marcksl1a*$^{rk23}$;*marcksl1b*$^{rk24}$ mutants at 30 and 52 hpf. Scale bar, 250 µm.

**Supplementary Fig. 5. Marcksl1-induced increase in vessel diameter and membrane blebbing occur in the presence of pericytes.**

**a – c**, Maximum intensity projection of a confocal z-stack of an ISV and DLAV of *TgBAC(pdgfrb:GFP)*$^{ncv22}$ with mosaic expression of marcksl1b-T2A-mKate2CAAX in ECs (magenta) at 54 hpf. Pericytes are labelled green. *, regions of vessel that are wider. **b** and **c**, Still images of a time-lapse movie. Arrow, basal blebs protrude in areas of blood vessel wrapped by pericytes. 00:00, hh:mm. Scale bars, 20 µm (a) and 5 µm (b and c).

**Supplementary Fig 6. Marcksl1 promotes endothelial cell proliferation.**

**a**, Maximum intensity projection of confocal z-stacks of ISVs of 52 hpf wildtype and *marcksl1* mutant embryos in *Tg(fli1ep:Lifeact-EGFP)*$^{zf495}$;*Tg(kdr-l:ras-mCherry)*$^{s916}$ background. Cell nuclei are detected by DAPI staining. Arrows indicate endothelial nuclei in ISVs. Scale bar, 20 µm. **b - c**, Quantification of EC number in arterial (**b**) and venous (**c**) ISVs at 52 hpf. Graphs show the relative frequency of nucleus number found in each ISV (wildtype: *n*=139 aISVs/111 vISVs from 21 embryos; *marcksl1a*$^{rk23}$: *n*=121 aISVs/163 vISVs from 20 embryos; *marcksl1b*$^{rk24}$: *n*=156 aISVs/124 vISVs from 20 embryos; *marcksl1a*$^{rk23}$;*marcksl1b*$^{rk24}$: *n*=167 aISVs/86 vISVs from 18 *marcksl1a*$^{rk23}$;*marcksl1b*$^{rk24}$ embryos). Data are collected from 2 independent experiments. **d - e**, Quantification of EC divisions in arterial (**d**) and venous (**e**) ISVs from time-lapse movies from 24 hpf to 48 hpf. Graphs show the relative frequency of cell divisions found in ISVs (wildtype: *n*=31 aISVs/11 vISVs from 11 embryos; *marcksl1a*$^{rk23}$;*marcksl1b*$^{rk24}$: *n*=23 aISVs/17 vISVs from 9 embryos). Data are collected from 3 independent experiments. **f**, Quantification of ISVs with one pH3 positive EC in wildtype and *marcksl1a*$^{rk23}$;*marcksl1b*$^{rk24}$ embryos at 30 hpf (*n*=30 wildtype/30 mutant embryos), 36 hpf (n=30 wildtype/26 mutant embryos) and 48 hpf (*n*=23 wildtype/32 mutant embryos). Statistical significance was determined by two-tailed Student's *t* test. $^{*}P < 0.05$, $^{**}P < 0.01$, ns, not significant. Mean ± S. D.

**Supplementary Fig. 7. Knockdown of *MARCKSL1* decreases HUVEC size.**

**a**, *MARCKSL1* mRNA expression level after siRNA knockdown (*n*=3 independent experiments). **b**, Maximum intensity projection of confocal z-stacks of HUVECs transfected with control non-targeting siRNA or MARCKSL1 siRNA 2 days post-transfection. **c – e**, Quantification of cell area (**c**), cell spikiness index (**d**) and cell aspect ratio (**e**; siCONTROL: *n*=233 cells; siMARCKSL1: *n*=431 cells from 3 independent experiments). Statistical significance was assessed by ordinary one-way ANOVA with Mann-Whitney *U* test. $^{*}P < 0.05$, $^{**}P < 0.01$, $^{***}P < 0.001$, $^{****}P < 0.0001$. Mean values are indicated. Scale bars, 50 µm.

**Supplementary Fig. 8. Marcksl1 induces ectopic blebbing in hindbrain vessels.**

Maximum intensity projection of a confocal z-stack of hindbrain vessels of 72 hpf *Tg(fli1:Lifeact-mCherry)$^{ncv7}$* embryos with mosaic expression of Marcksl1b-EGFP in ECs. Arrows, local dilation of blood vessels. Arrowheads, blebs. Scale bar, 50 µm.

**Supplementary Fig. 9. Methodology to quantify membrane blebbing.**

Membrane blebbing index is measured as a ratio between membrane length of the bleb and Euclidean length at each timepoint of a times series. The spectral power density was then obtained and averaged over a window of interest (0.04 – 0.06 Hz) during which blebs form. The average spectral power density for wildtype and Marcksl1-overexpressing (OE) ECs was plotted.

**Supplementary Fig. 10. *MARCKSL1* mRNA expression is not regulated by shear stress *in vitro*.**

HPAECs were exposed to static or 15 dyn cm$^{-2}$ laminar shear stress for 0.5, 1 or 6 hours. *MARCKSL1* mRNA expression is shown relative to *GAPDH* expression. Data are mean ± s.d. of 3 independent experiments. Statistical analysis was performed by one-way ANOVA and Turkey's multiple comparison test. ns, not significant.

**Supplementary Fig. 11. JNK phosphorylation of Marcksl1 alters EC shape.**

Maximum intensity projections of confocal z-stacks of HUVECs transfected with dephospho-Marcksl1-EGFP (**a**, Marcksl1-AAA-EGFP) and phosphomimetic Marcksl1-EGFP (**b**, Marcksl1-DDD-EGFP) and stained with DAPI (blue) and phalloidin (red). Quantification of cell area (**c**), cell spikiness index (**d**) and cell aspect ratio (**e**; Marcksl1-AAA-EGFP: *n*=210 cells; Marcksl1-DDD-EGFP: *n*=200 cells). Analyzed by unpaired Student's *t*-test. ns, not significant. Scale bars, 25 µm.

**Supplementary Fig. 12. Increased ISV regression in *marcksl1a$^{rk23}$;marcksl1b$^{rk24}$* embryos**

**a**, Maximum intensity projection of a confocal z-stack of ISVs in zebrafish trunk of 2 dpf *marcksl1a$^{rk23}$;marcksl1b$^{rk24}$* embryo in *Tg(fli1ep:Lifeact-EGFP)$^{zf495}$;Tg(kdr-l:ras-mCherry)$^{s916}$* background. **b**, Relative frequency of vessel phenotypes in wildtype and *marcksl1* mutant embryos (wildtype: $n$=45 ISVs/11 embryos; *marcksl1a*$^{rk23}$: $n$=116 ISVs/29 embryos; *marcksl1b*$^{rk24}$: $n$=100 ISVs/25 embryos; *marcksl1a*$^{rk23}$;*marcksl1b*$^{rk24}$: $n$=126 ISVs/30 embryos). Scale bar, 20 µm.

**Supplementary Fig. 13. *Marcksl1a* and *marcksl1b* mutant alleles**

CRISPR/Cas9-induced mutations in *marcksl1a* gene and TALEN-induced mutations in *marcksl1b* gene. Partial genomic and protein (MH2 and Effector Domain are highlighted) sequences are shown. Missense amino acids and nucleotide substitutions/insertions are shown red.

Six *marcksl1a* mutants were obtained (**a**). Mutant alleles from 1 to 3 had a 3-nt, 18-nt and 21-nt deletions leading to 1, 6 and 7 amino acids deletions from protein, respectively, without affecting the open reading frame (ORF). Mutant allele 5 had a 7-nt deletion leading to a frameshift after T51 and premature stop codon at amino acid 94 after 43 missense amino acids. Mutant allele 6 had an 8-nt deletion leading to a frameshift after G50 and premature stop codon at amino acid 54 after 4 missense amino acids. Mutant allele ***rk23*** (used in all experiments) has a 2-nt deletion which leads to a frameshift after T51 and premature stop codon at amino acid 56 after 5 missense amino acids.

Five *marcksl1b* mutants were obtained (**b**). Mutant alleles 1 and 2 had a 3-nt and 18-nt deletions, respectively, leading to 1 and 6 amino acids deletions from protein without affecting the ORF. Mutant allele 4 had a 2-nt insertion leading to a frameshift after E13 and premature stop codon at amino acid 100 after 87 missense amino acids. Mutant allele 5 had a 1-nt insertion leading to a frameshift after G14 and premature stop codon at amino acid 19 after 5 missense amino acids. Mutant allele ***rk24*** (used in all experiments) has a 5-nt deletion which leads to a frameshift after E13 and premature stop codon at amino acid 17 after 4 missense amino acids.

#### SUPPLEMENTARY TABLES

**Supplementary Table 1. Plasmids used in this study**

**Supplementary Table 2. Oligonucleotides used in this study**

#### SUPPLEMENTARY VIDEO LEGENDS

**Supplementary video 1. Formation of ISVs and DLAV in wildtype and *marcksl1a*$^{rk23}$*;* *marcksl1b*$^{rk24}$ embryos.**

Live-imaging of wildtype and mutant embryos in *Tg(fli1ep:Lifeact-EGFP)*$^{zf495}$*;Tg(kdr-l:ras-mCherry)*$^{s916}$ background. 00:00, hours:minutes post fertilization.

**Supplementary video 2. Marcksl1 overexpression deregulates blood vessel diameter.**

Left panel: time-lapse imaging of an ISV of a 2 dpf *Tg(kdr-l:ras-mCherry)*$^{s916}$ embryo. Right panel: an ISV with mosaic Marcksl1b-EGFP overexpression (OE) at 2 dpf. Increased Marcksl1b level leads to fluctuation in vessel diameter as well as ectopic filopodia formation and blebbing. 00:00, hours:minutes.

**Supplementary video 3. Ectopic Marcksl1a expression in ECs induces excessive filopodia formation and membrane blebbing during ISV lumenization.**

ECs expressing Marcksl1a-T2A-mKate2CAAX are in magenta while wildtype ECs are green. Movie was taken from a 31 hpf *Tg(fli1ep:Lifeact-EGFP)*$^{zf495}$ embryo. 00:00; hours:minutes.

**Supplementary video 4. Ectopic Marcksl1b expression in ECs induces basal blebbing in perfused vessels.**

ECs expressing Marcksl1b-EGFP are in magenta while wildtype ECs are in green. Movie was taken from a 54 hpf *Tg(fli1:Lifeact-mCherry)*$^{ncv7}$ embryo. For wildtype EC membrane behaviour, refer to control in Supplementary Video 2. 00:00; hours:minutes.

**Supplementary video 5. Marcksl1-induced blebs are filled with blood plasma.**

Time-lapse imaging of DLAV of 2 dpf Tg(*Tg(kdr-l:ras-mCherry)*$^{s916}$ embryo (wildtype) and embryo with endothelial overexpression of Marcksl1-EGFP. Lumen is labelled with Dextran-Rhodamine (magenta) while endothelial plasma membrane is in green. Marcksl1-induced blebs are comprised of apical and basal membranes (green). 00:00; minutes:seconds.

**Supplementary video 6. Actin dynamics in Marcksl1-induced blebs.**

2 dpf *Tg(fli1:Lifeact-mCherry)$^{ncv7}$* embryo with mosaic expression of Marcksl1b-EGFP. EC with increased Marcksl1b expression (magenta) exhibit increased filopodia formation, basal blebs and irregular cell shape. In most blebs, actin (green) reassembly around the bleb cortex precedes retraction (box A) while failure to do so leads to bleb persistence (box B). Refer to Supplementary Video 11 for actin dynamics in a control embryo. 00:00; minutes:seconds.

**Supplementary video 7. Non-muscle myosin II dynamics in Marcksl1-induced blebs.**

Wildtype: dynamics of Myl9b-EGFP (green) in an ISV of a 2 dpf *Tg(fli1ep:myl9b-EGFP)$^{rk25}$;Tg(kdr-l:ras-mCherry)$^{s916}$* control embryo. Endothelial membrane is in magenta. Marcksl1 overexpression: 2 dpf *Tg(fli1ep:myl9b-EGFP)$^{rk25}$* embryo with mosaic overexpression of Marcksl1b-EGFP. Myl9b-EGFP (green) accumulates at the neck of the bleb and eventually reassembles at the front of the bleb (magenta) before retraction. 00:00; minutes:seconds.

**Supplementary video 8. Local weakening of EC cortex induces local membrane blebbing.**

Laser ablation of an arterial ISV was performed on a 3 dpf *Tg(kdr-l:ras-mCherry)$^{s916}$* embryo. *, site of ablation. Note that surrounding membranes not ablated by laser remain unperturbed. Time-lapse imaging at a single plane of the vessel was taken. 00:00:00, minutes:seconds:milliseconds.

**Supplementary video 9. Short-term inhibition of actin polymerization induces membrane blebbing in perfused blood vessels.**

2 dpf *Tg(kdr-l:ras-mCherry)$^{s916}$* embryos were treated with 0.4% DMSO (left panel) or 0.3 µg/ml Latrunculin B (right panel) and imaged 10 minutes later. Magenta arrow, apical bleb. Black arrow, basal bleb. 00:00, hours:minutes.

**Supplementary video 10. Decreased blood flow normalizes Marcksl1-induced blebbing.**

2 dpf *Tg(fli1:Lifeact-mCherry)*$^{ncv7}$ embryo with mosaic expression of Marcksl1b-EGFP was treated with 1X tricaine for 2 hours, 4X tricaine for 4 hours, washed for 1 hour and then returned to 1X tricaine for 3 hours. Magenta, Marcksl1b-overexpressing cell; green, actin. 00:00, hours:minutes after start of each treatment.

**Supplementary video 11. Cortical actomyosin network is highly dynamic.**

Time-lapse imaging of a perfused ISV from a 2 dpf *Tg(fli1ep:myl9b-EGFP)*$^{rk25}$;*Tg(fli1:Lifeact-mCherry)*$^{ncv7}$ embryo. The EC cortex is composed of a dynamic meshwork of actin (magenta, top panel) and myosin II (green, bottom panel). 00:00; minutes:seconds.

**Supplementary video 12. Ectopic expression of Fascin1a in ECs induces bleb formation and vessel dilation in perfused blood vessels.**

Time-lapse imaging of lumen formation in ISVs and DLAV of *Tg(fli1ep:Lifeact-EGFP)*$^{zf495}$ embryo with mosaic expression of Fascin1a-T2A-mKate2CAAX from 30 hpf. Magenta, EC with Fascin1a overexpression; green, actin/wildtype ECs. 00:00; hours:minutes.

**Supplementary video 13. Inhibition of Arp2/3-mediated branched actin formation leads to membrane blebbing and deregulation of vessel diameter.**

2 dpf *Tg(kdr-l:ras-mCherry)*$^{s916}$ embryos were treated with 0.4% DMSO or 200µM CK666 for 1 hour and then imaged. 00:00, minutes:seconds.

Supplementary Figure 1

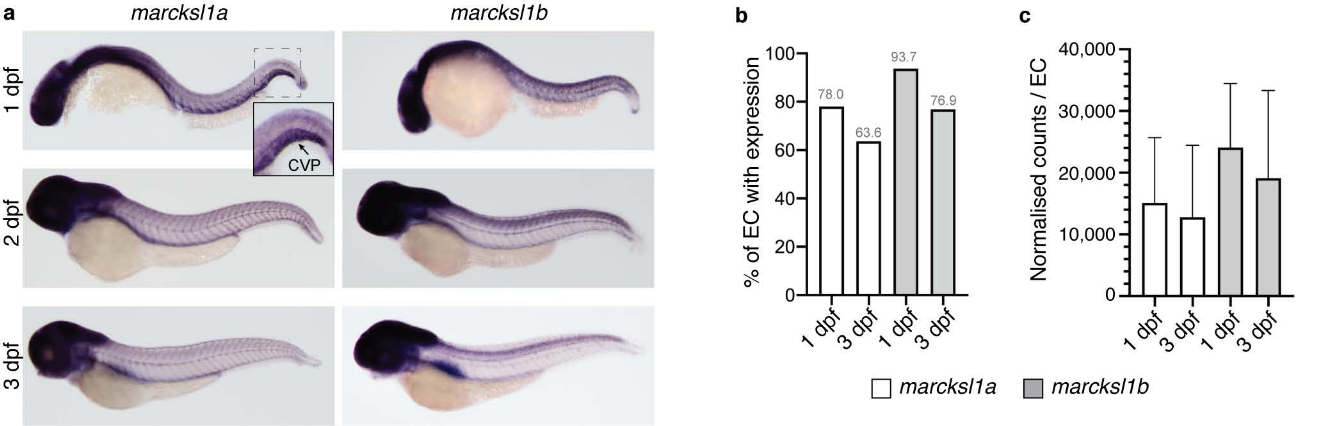

### Supplementary Figure 2

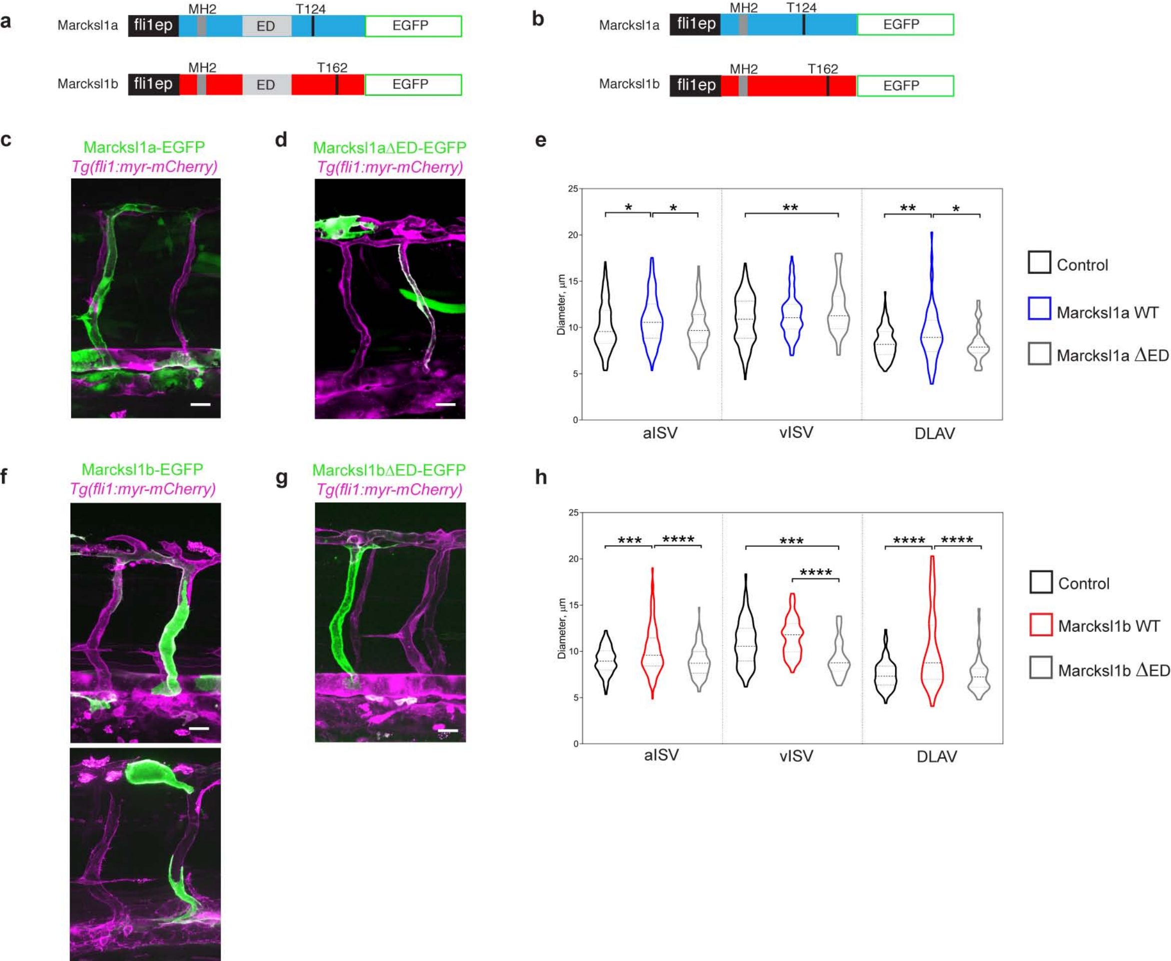

### Supplementary Figure 3

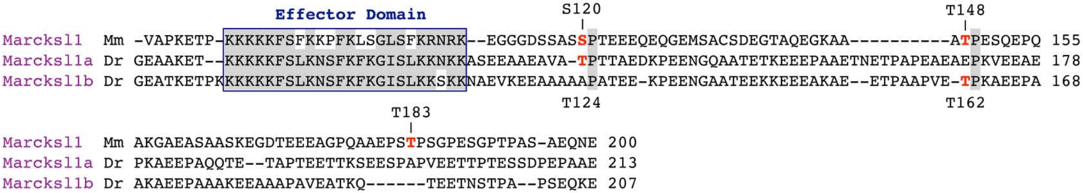

Supplementary Figure 4

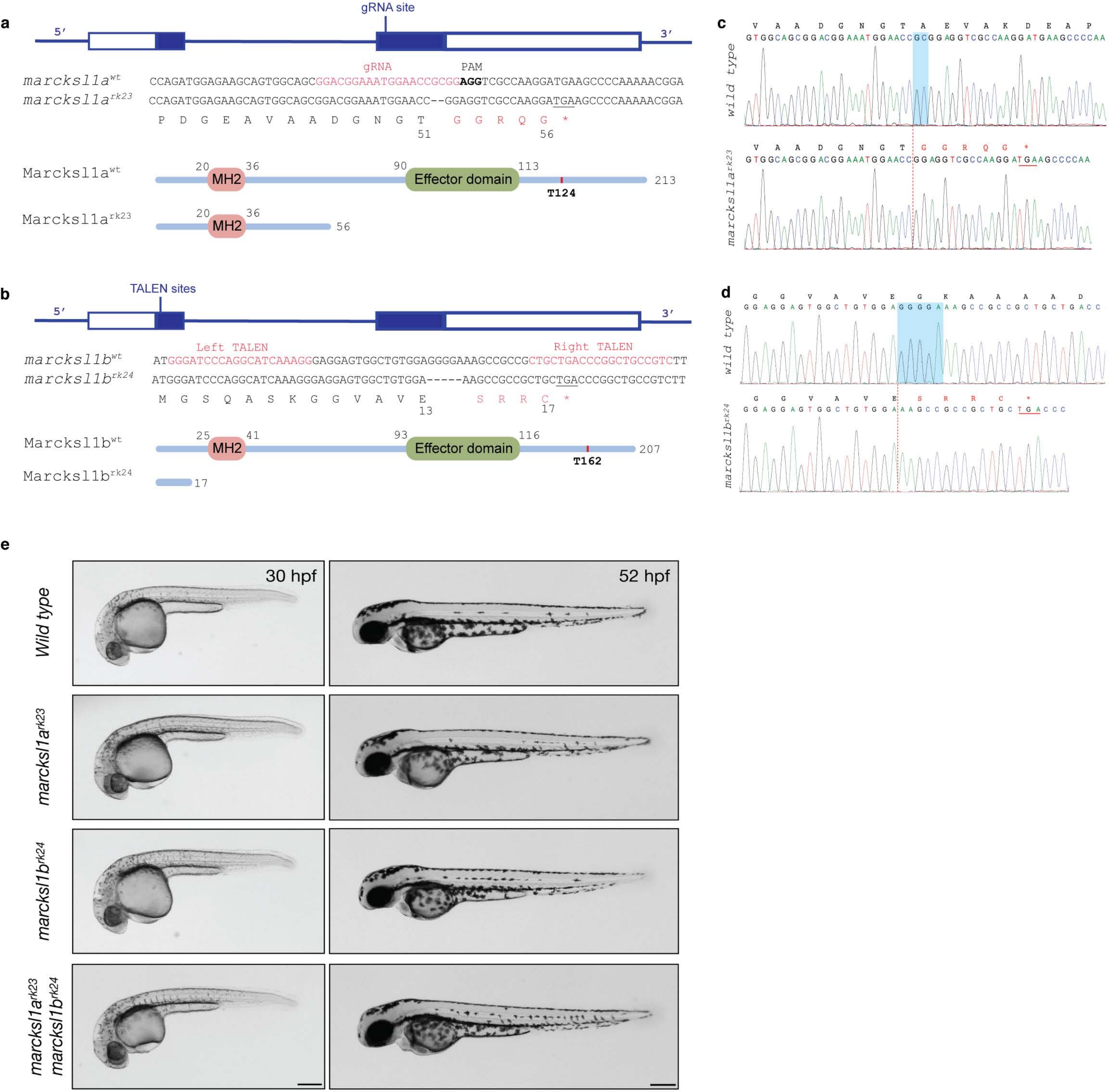

Supplementary Figure 5

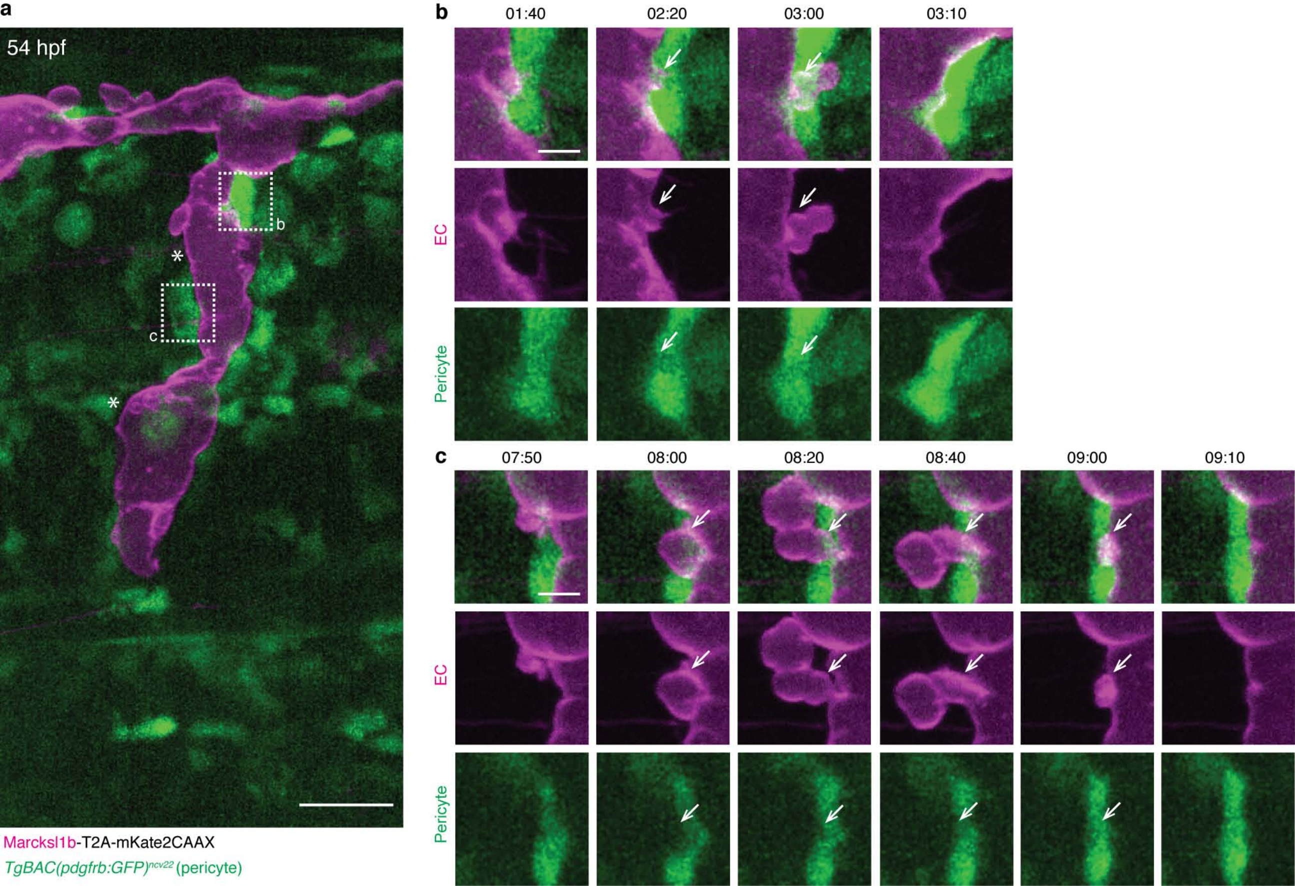

### Supplementary Figure 6

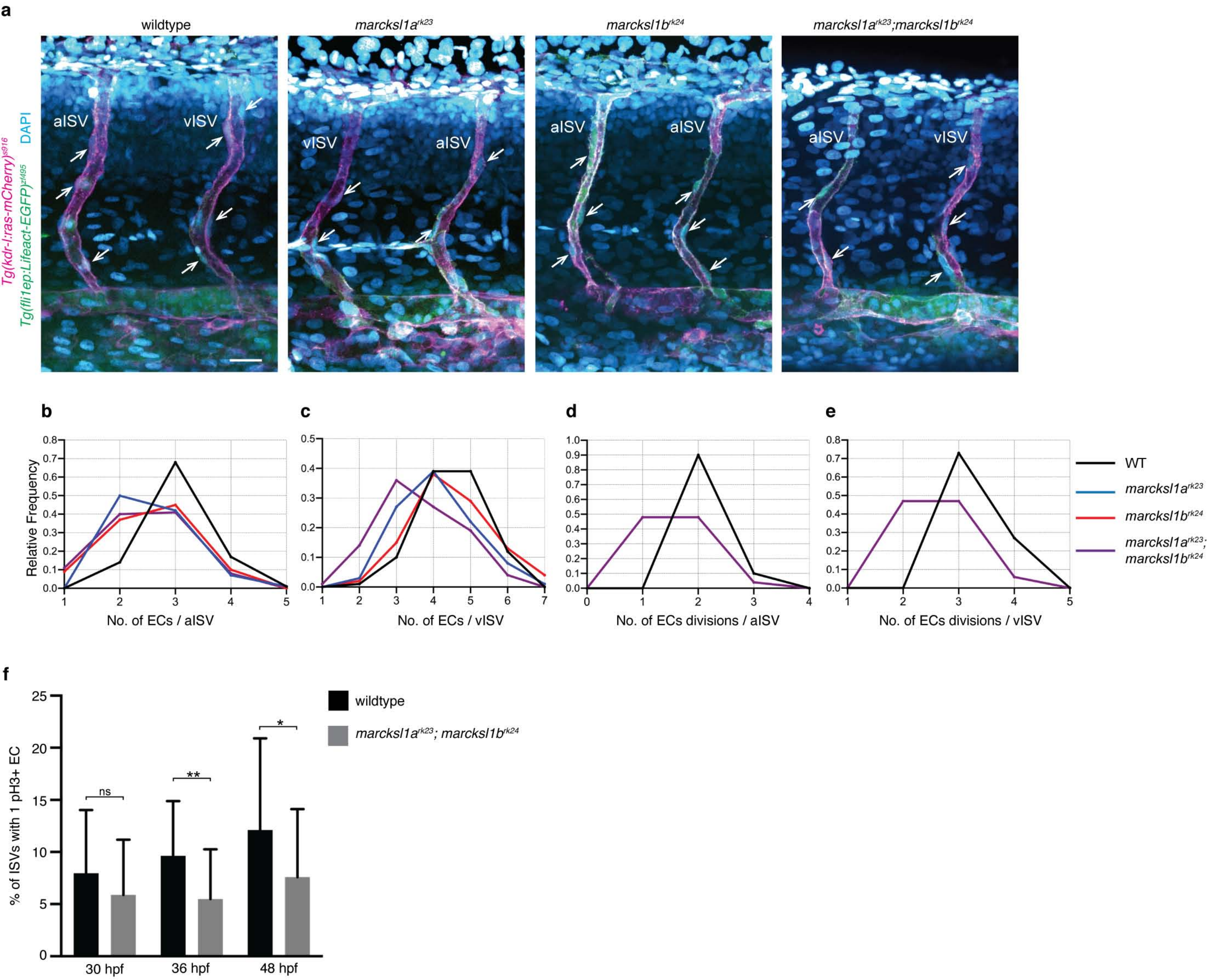

### Supplementary Figure 7

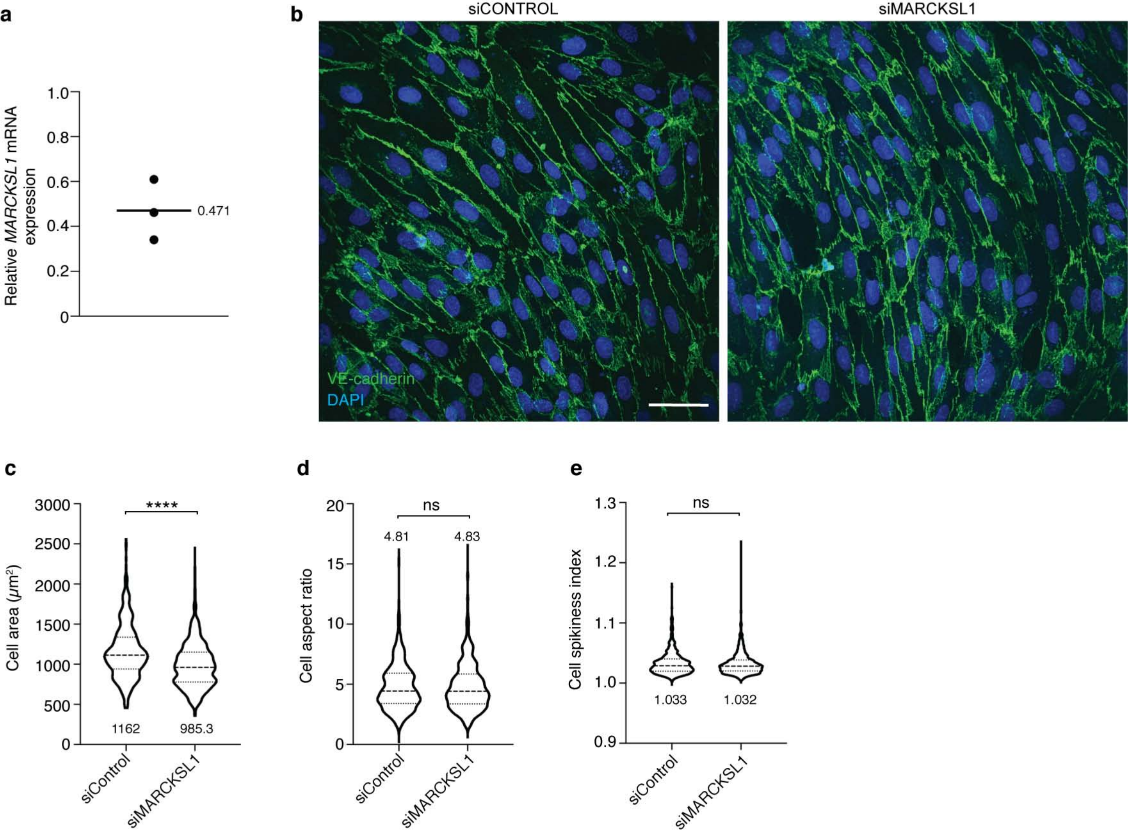

Supplementary Figure 8

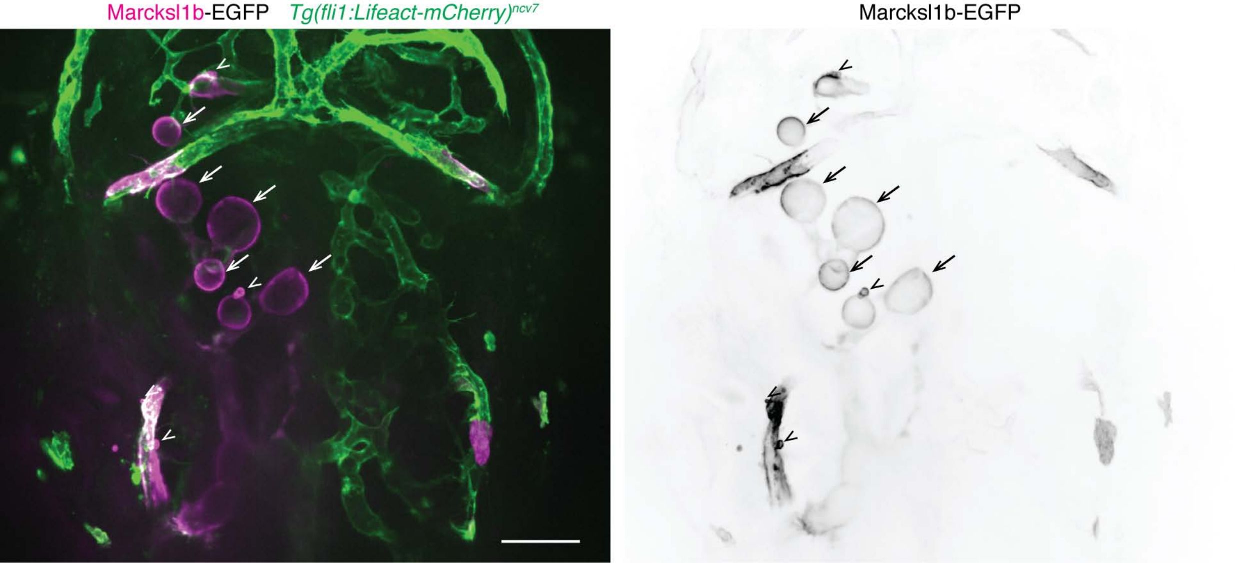

### Supplementary Figure 9

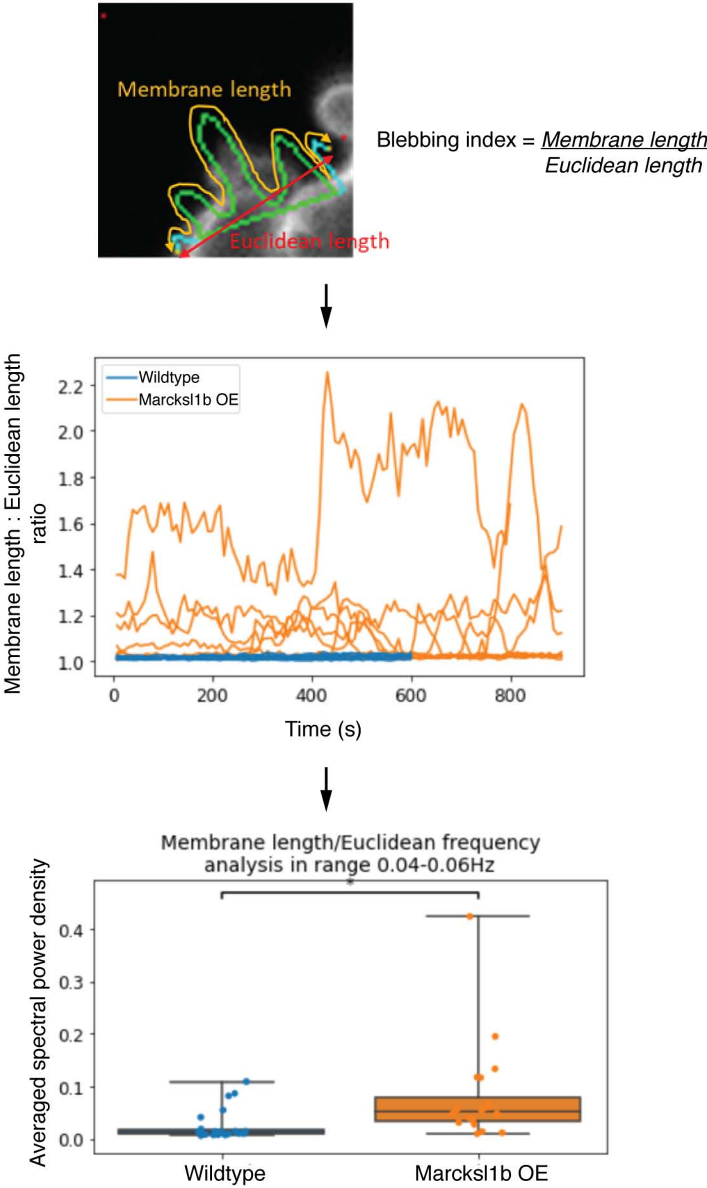

### Supplementary Figure 10

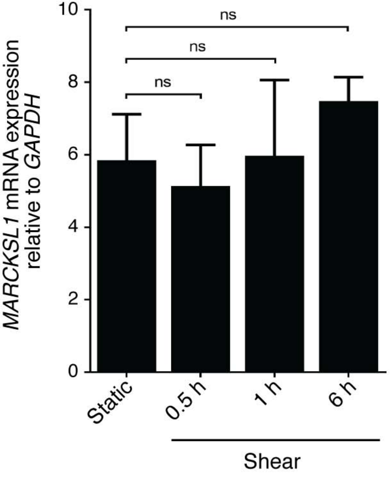

### Supplementary Figure 11

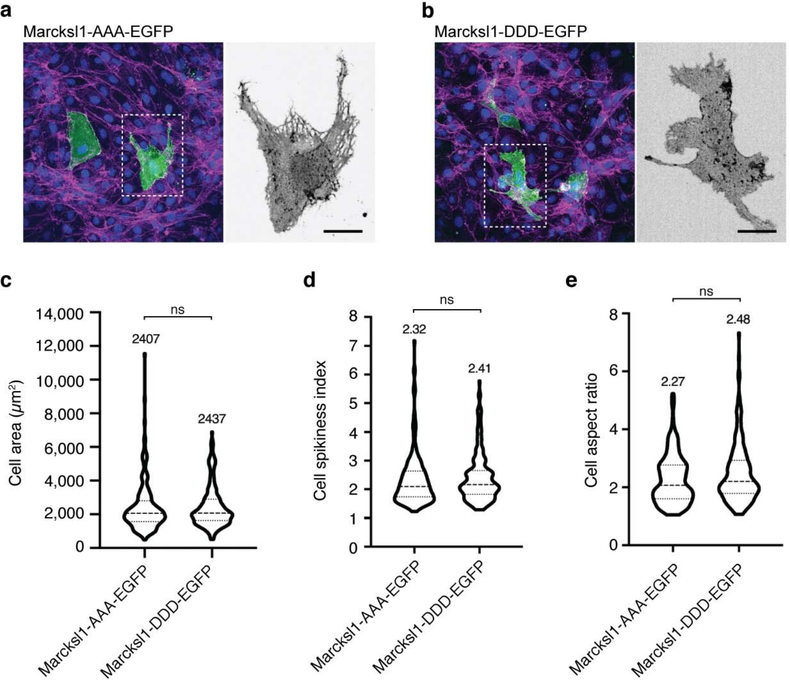

Supplementary Figure 12

a

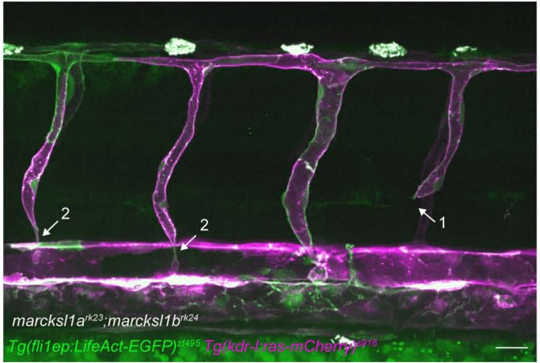


b

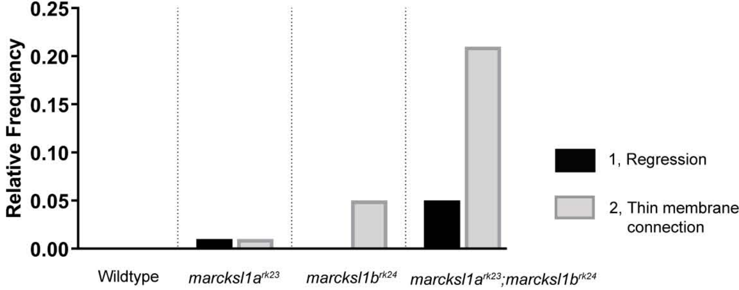

Supplementary Figure 13

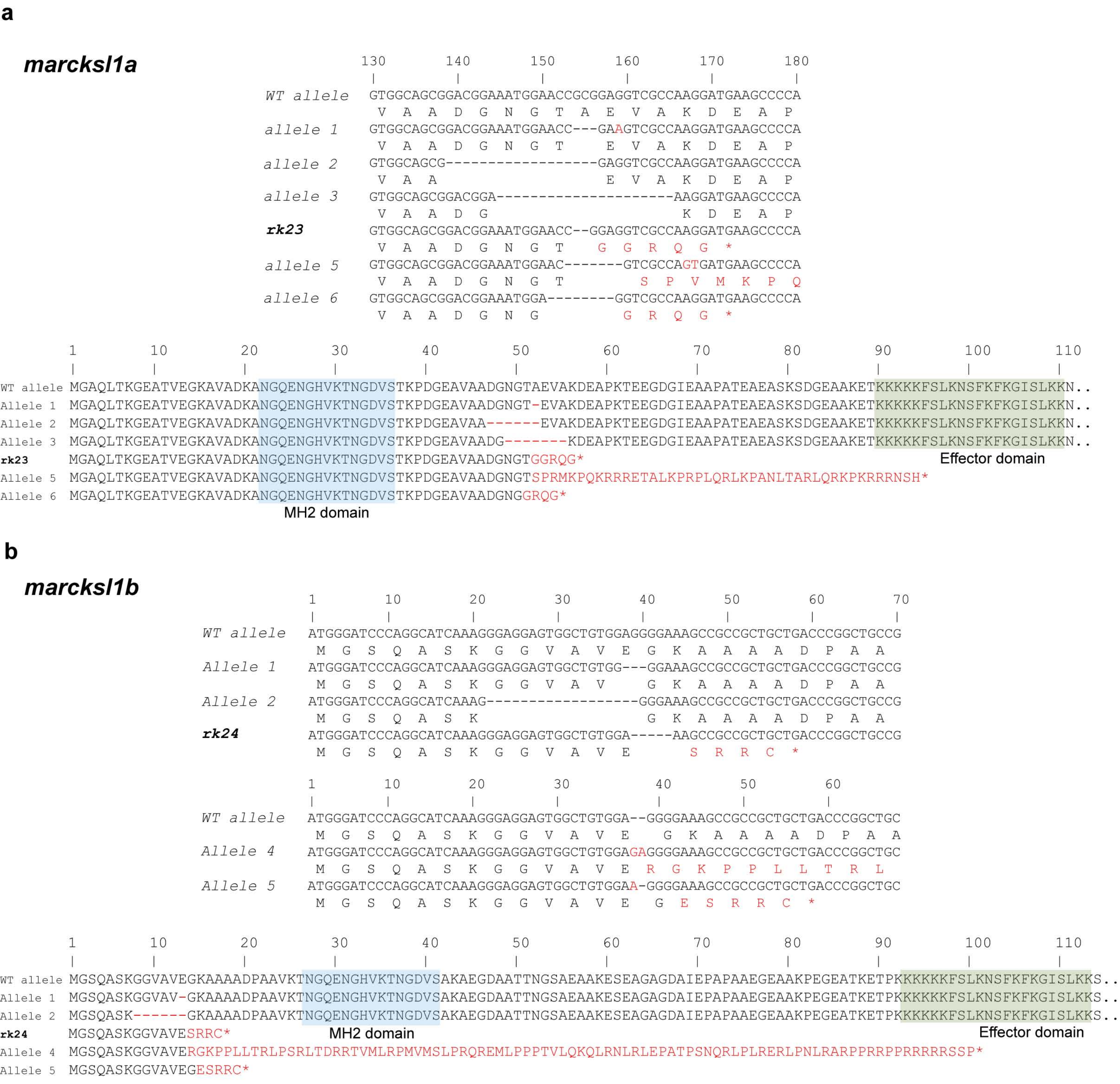

**Supplementary Table 1. Plasmids used in this study**

| Plasmid name | Plasmid backbone | Method of generation | Reference | Experiments | Comments |
|---|---|---|---|---|---|
| *pMiniT-marcksl1a* | pMiniT 2.0 (NEB) | PCR cloning | This paper | Suppl.Fig.1a | For riboprobe synthesis (1127 nt including 300 nt of 5'UTR and 185 nt of 3'UTR) |
| *pMiniT-marcksl1b* | pMiniT 2.0 (NEB) | PCR cloning | This paper | Suppl.Fig.1a | For riboprobe synthesis (1125 nt including 217 nt of 5'UTR and 284 nt of 3'UTR) |
| *pMiniT-fscn1a* | pMiniT 2.0 (NEB) | PCR cloning | This paper | | Used to construct *6xUAS:fscn1a-T2A-mKate2CAAX* plasmid |
| **Expression constructs** | | | | | |
| *fli1ep:marcksl1a-EGFP* | pDestTol2CG2-*cmlc2:EGFP* | Gateway cloning | This paper | Suppl.Fig.2 | Mosaic overexpression of wildtype Marcksl1a in ECs |
| *fli1ep:marcksl1aΔED-EGFP* | pDestTol2CG2-*cmlc2:EGFP* | Site-Directed mutagenesis | This paper | | Mosaic overexpression of mutated (without ED) Marcksl1a in ECs |
| *fli1ep:marcksl1b-EGFP* | pDestTol2CG2-*cmlc2:EGFP* | In-Fusion cloning | This paper | | Mosaic overexpression of wildtype Marcksl1b in ECs |
| *fli1ep:marcksl1bΔED-EGFP* | pDestTol2CG2-*cmlc2:EGFP* | Site-Directed mutagenesis | This paper | | Mosaic overexpression of mutated (without ED) Marcksl1b in ECs |
| *6xUAS:marcksl1a-T2A-mKate2CAAX* | pDestTol2CG2-*cry:mKate2* | In-Fusion cloning | This paper | Fig.2a-c, 5a,c | Mosaic overexpression of wildtype Marcksl1a in ECs |
| *6xUAS:marcksl1aΔED-T2A-mKate2-CAAX* | pDestTol2CG2-*cry:mKate2* | Site-Directed mutagenesis | This paper | Fig.2a-c | Mosaic overexpression of mutated (without ED) Marcksl1a in ECs |
| *6xUAS:marcksl1aT124A-T2A-mKate2-CAAX* | pDestTol2CG2-*cry:mKate2* | Site-Directed mutagenesis | This paper | | Mosaic overexpression of dephospho-Marcksl1a in ECs |
| *6xUAS:marcksl1aT124D-T2A-mKate2-CAAX* | pDestTol2CG2-*cry:mKate2* | Site-Directed mutagenesis | This paper | | Mosaic overexpression of phosphomimetic Marcksl1a in ECs |
| *6xUAS:marcksl1b-T2A-mKate2CAAX* | pDestTol2CG2-*cry:mKate2* | In-Fusion cloning | This paper | Fig.2d-f, 4a,c,d, 5b,d-f, 6, Suppl.Fig.5,8 | Mosaic overexpression of wildtype Marcksl1b in ECs |
| *6xUAS:marcksl1bΔED-T2A-mKate2-CAAX* | pDestTol2CG2-*cry:mKate2* | Site-Directed mutagenesis | This paper | Fig.2d-f | Mosaic overexpression of mutated (without ED) Marcksl1b in ECs |
| *6xUAS:marcksl1bT162A-T2A-mKate2-CAAX* | pDestTol2CG2-*cry:mKate2* | Site-Directed mutagenesis | This paper | | Mosaic overexpression of dephospho-Marcksl1b in ECs |
| *6xUAS:marcksl1bT162D-T2A-mKate2-CAAX* | pDestTol2CG2-*cry:mKate2* | Site-Directed mutagenesis | This paper | | Mosaic overexpression of phosphomimetic Marcksl1b in ECs |
| *6xUAS:fscn1a-T2A-mKate2CAAX* | pDestTol2CG2-*cry:mKate2* | In-Fusion cloning | This paper | Fig.9a | Mosaic overexpression in ECs |
| *fli1ep:lynEGFP* | pDestTol2CG2-*cmlc2:EGFP* | Gateway cloning | This paper | Fig.4c,d | *In vivo* cell shape analysis (as a control) |
| *fli1ep:Myl9b-EGFP* | pDestTol2CG2-*cmlc2:EGFP* | Gateway cloning | This paper | Fig.1d-f, 6c, 8c | Used to generate *Tg(fli1ep:Myl9b-EGFP)* transgenic line |
| *fli1ep:EGFP-PLCd1PH* | pDestTol2CG2-*cmlc2:EGFP* | Gateway cloning | This paper | Fig.1a | Used to generate *Tg(fli1ep:EGFP-PLCd1PH)* transgenic line |
| *pEGFP-N1* | | | Clontech | Fig.4f-i, 8e-g | *In vitro* cell shape analysis and actin organization (as a control) |
| *pEGFP-U6* | pEGFP-N1 | | PMID: 18840295 | | Used to construct shRNA vplasmids |
| **shRNA plasmids** | | | | | |
| *pEGFP-CAAX-U6:shMARCKSL1*<br>*pEGFP-CAAX-U6:shControl* | pEGFP-U6 | Restriction/ligation cloning | This paper | MARCKSL1 KD (*in vitro* cell shape analysis and actin organization) Fig.4j-m, 8h-i | Original pEGFP-U6 vector was modified by in-frame fusion of the last 21 amino acids of human H-ras (CAAX box) to the C-terminus of EGFP, creating a membrane-targed EGFP. |
| **mouse Marcksl1 plasmids** | | | | | |
| *pMarcksl1-EGFP* | pEGFP-N1 | | PMID: 22751924 | Fig.4f-i, 8d-g | *In vitro* cell shape analysis, actin organization |
| *pMarcksl1-AAA-EGFP* | | | | Fig.8e-g, Suppl.Fig.11 | *In vitro* cell shape analysis (dephospho-Marcksl1), actin organization |
| *pMarcksl1-DDD-EGFP* | | | | Fig.8e-g, Suppl.Fig.11 | *In vitro* cell shape analysis (phosphomimetic Marcksl1), actin organization |
| *p5E-fli1ep* | | | Gift from Nathan Lawson (Univ. Massachusetts Medical School) | | Used as a source of *fli1ep* promotor |
| *pDestTol2-cry:mKate2* | | | Gift from Darren Gilmour (EMBL, Heidelberg) | | Used as a backbone plasmid for In-Fusion cloning |
| *pME-lynEGFP* | | | Gift from Darren Gilmour (EMBL, Heidelberg) | | To construct *fli1ep:lynEGFP* plasmid |
| *pME-Myl9b* | | | Gift from Holger Gerhardt (MDC, Berlin) | | To construct *fli1ep:Myl9b-EGFP* plasmid |
| *PLCd1PH* | | | Gift from Carsten Schultz (EMBL, Heidelberg) | | To construct *fli1ep:EGFP-PLCd1PH* plasmid |

#### Supplementary Table 2. Oligonucleotides used in this study.

All synthetic oligonucleotides were purchased from Fasmac (Japan) and Invitrogen

**Primers for genotyping**

| | |
|---|---|
| Marcksl1a-Fwd | GTGTGTGTGTTTGCCAAGATGCATT |
| Marcksl1a-Rev | GGTTGCCACAGCGATATGAGATCAC |
| Marcksl1b-Fwd | AGCGCTGTAGGACTGGAACTGGTA |
| Marcksl1b-Rev | CAACGACAGAAATGAATCGAAAACG |

**Primers for the full-length cDNA cloning**

| | |
|---|---|
| marcksl1a-fwd | ATGAAGCTCCAGCCCTCTTGTGCAGA |
| marcksl1a-rev | CGCGAACCAGTGAACGTTATCAGCA |
| marcksl1b-fwd | AGCGCTGTAGGACTGGAACTGGTA |
| marcksl1b-rev | TCCTCAACTCACTCTTGTGCTGACA |
| fscn1a-fwd | CATCATCCACGGTGCACCAGCAGA |
| fscn1a-rev | TGGCCACTCGTCAGGTCATCGA |

**Primers for mutagenesis (deletion of an Effector domain)**

| | |
|---|---|
| marcksl1a-delED-fwd | GCAAGTGAGGAGGCAGCGGA |
| marcksl1a-delED-rev | GGTTTCCTTTGCAGCCTCGC |
| marcksl1b-delED-fwd | AATGCTGAGGTGAAGGAAGAGGC |
| marcksl1b-delED-rev | CTTGGGGGTCTCCTTGGTGG |

**Primers for mutagenesis (T124A, T124D)**

| | |
|---|---|
| marcksl1aT124A-fwd | GGCTGTGGCCgcaCCCACCACCG |
| marcksl1aT124D-fwd | GGCTGTGGCCgacCCCACCACCGC |
| marcksl1aT124-rev | TCCGCTGCCTCCTCACTTGC |

**Primers for mutagenesis (T162A, T162D)**

| | |
|---|---|
| marcksl1bT162A-fwd | CCCTGTTGAAgCCCCCAAGGC |
| marcksl1bT162D-fwd | CCCTGTTGAAgaCCCCAAGGCCGAGG |
| marcksl1bT162-rev | GCAGCGGGTGTCTCCTCG |

**Oligos for human MARCKSL1 shRNA construction**

| | |
|---|---|
| shRNA sense | GTGTGAACGGAACAGATGATG CCTGACCCACATCATCTGTTCCGTTCACACTTTTTTG |
| shRNA antisense | AATTCAAAAAAGTGTGAACGGAACAGATGATGTGGGTCAGGGATCATCTGTTCCGTTCACAC |
| shRNA scrambled sense | GAGTCATGGGTCAGTTATATGCCTGACCCACATATAACTGACCCATGACTCTTTTTTG |
| shRNA srambled antisense | AATTCAAAAAAGAGTCATGGGTCAGTTATATGTGGGTCAGGCATATAACTGACCCATGACTC |

**Oligos for marcksl1a sgRNA construction**

| | |
|---|---|
| tracrRNA | AAAAGCACCGACTCGGTGCCACTTTTTCAAGTTGATAACGGACTAGCCTTATTTTAACTTGCTATTTCTAGCTCTAAAAC |
| sgRNA marcksl1a | TAATACGACTCACTATAGGACGGAAATGGAACCGCGG GTTTTAGAGCTAGAAATAGCAAG |

**qPCR primers**

| | |
|---|---|
| MARCKSL1-fwd | ATCATGGGCAGCCAGAGCT |
| MARCKSL1-rev | TGCCTCATCTGTTCCGTTCACAG |
| GAPDH-fwd | GCCACATCGCTCAGACACCAT |
| GAPDH-rev | TGAAGGGGTCATTGATGGCAACA |
